# SNP genotyping and population analysis of five indigenous Kazakh sheep breeds

**DOI:** 10.1101/2020.04.28.065797

**Authors:** Alexandr Pozharskiy, Aigerim Khamzina, Dilyara Gritsenko, Zhangylsyn Khamzina, Shinara Kassymbekova, Nurlybek Karimov, Talgat Karymsakov, Nurlan Tlevlesov

## Abstract

Sheep husbandry is an important branch of agriculture in Kazakhstan. Modern agrarian and breeding science demands inclusion of molecular genetic and genomic data to supplement traditional methods. Here we used medium-scale SNP genotyping for the first time to determine the population structure of five local sheep breeds in Kazakhstan and their relation to global sheep diversity.

Principal component analysis and model-based structure analysis of general population markers revealed two breed groups. The first group included Akzhayik and Kazak Fine-wool sheep and the second group had Edilbay, Saryarka and Kazakh Semi-coarse wool sheep. High heterogeneity of different populations of Akzhayik and Kazakh Semi-coarse wool sheep was observed. A neighborjoining tree comparing Kazakh sheep data with the dataset generated by the Sheep HapMap project supported a close relationship between Kazakh sheep varieties and ancient domestic sheep ancestors.

## Introduction

Domestic sheep (*Ovis aries* Linnaeus) are one of the most culturally and economically important farm animals. Sheep were domesticated between 11,000 and 9,000 B.C. in Central and Southwestern Asia (Tapio et al., 2010; Zeder, 2008). In Kazakhstan, sheep together with horses are important domestic animals and have a long history of traditional husbandry. According to archaeological data, domestic sheep were present in regions of Kazakhstan since the early bronze age (Frachetti and Benecke, 2009). Before the Soviet period, most Kazakh sheep varieties were the result of traditional folk breeding and selection. In the KazSSR, systematic work for the creation of new sheep breeds creation was carried out on state farms (Degen, 2013). Sheep husbandry is currently a key branch of the Kazakhstan economy and provides meat and wool for both domestic use and for export. Modern agriculture practices involve application of molecular genetic methods as a valuable tool to predict and select desirable traits and properties. The development of high throughput single-nucleotide polymorphism (SNP) genotyping technologies has contributed to the growing impact of genome-wide association studies (GWAS) in animal breeding, genomic prediction and animal selection (Gholizadeh et al., 2015).

SNPs are among the most common markers in the study of genetic processes. The advantages of SNPs are the high number of markers that can be processed simultaneously, the ability to cover both coding and non-coding regions, flexibility in genotyping of non-model organisms and low error rates across a range of DNA sample quality (Haynes and Latch, 2012). SNP genotyping is a powerful tool for marker-assisted selection in animal breeding (Seidel, 2010). With increased demand for meat and wool production by the sheep industry, breeders and geneticists are considering the genetic basis for interactions between genotype and environmental factors in order to maintain and improve the distinctive features and biological characteristics of various breeds (Zhang et al., 2013). One approach to improving breed features is identification of new gene pools that have potentially beneficial genotypes that can be introduced into the breeding process. From this perspective, indigenous sheep breeds in Kazakhstan are of particular interest, because, although they are well-established, few genomic studies have focused on these breeds.

In the present study we considered five sheep breeds that are widely used in Kazakhstan: Edilbay (Edilbayevskaya), Kazakh Fine-wool (Kazakh Fine-fleece), Akzhayik (Akzhayikskaya), Kazakh Semi-coarse wool sheep and Saryarka (Saryarkinskaya). The Edilbay breed is traditionally considered as a descendant of old sheep varieties in Kazakhstan. This breed was created at the end of the 19th century in the Ural region and is thought to have originated from breeding of Astrakhan sheep with local varieties that were naturally selected or the products of folk selection methods, although no documentary data concerning how this breed was established are available. Edilbay sheep are well adapted to the conditions of dry steppes, deserts, and graze nearly year-round (Ermekov and Golodnov, 1976). This breed is of special interest because it played a central role in creation of most local sheep breeds in Kazakhstan. Moreover, mitochondrial DNA analysis supports the genetic proximity of Edilbay to the hypothetical ancestor of modern sheep varieties (Hiendleder et al., 1998), although it has not been compared further with other sheep breeds that are common worldwide.

On September 28, 1945, the board of the USSR approved the Kazakh fine-wool breed of sheep, generated by V.A. Balmont, G.V. Bakanova and A.P. Pshenichny, as a new independent breed. Work on the breeding of this variety of fine-fleece sheep was begun by V.A. Balmont in regions of southeastern Kazakhstan where Kazakh fat tail queens were crossed with Prekos fine-wool sheep in the autumn of 1931. Animals resulting from this cross were characterized by robust growth, strong constitution and good adaptation to year-round pasture grazing (Balmont, 1957).

The Akzhayk meat-wool breed was bred in East Kazakhstan in 1996 using complex reproductive crossbreeding of fine-fleece and semi-fine-coarse-haired sheep with rams of the Lincoln and Romney march type and subsequent breeding of animals of the desired type “inside”. For many years, scientific guidance on selection and breeding of the Akzhayk breed was provided by V.V. Terentyev (Terentyev, 1987; Traisov et al., 2013).

The Saryarka fat tail breed by M.A. Ermekov et al. was approved in 1999. This breed of sheep was created over the period between 1970 and 1998 mainly through in-breeding selection and selection according to breeding traits of the improved Kazakh fat tail sheep with Edilbaev sheep as well as the partial blood line of the Kargaly semi-coarse sheep with subsequent inbreeding of animals of the desired type. These sheep are characterized by a strong constitution, a well-developed skeleton, regular physique, and strong limbs with a dense ungulate horn, which is important for year round pasture grazing. These sheep are found in Central Kazakhstan (Akhatov, 2006; Kanapin et al., 2001; Kanapin and Akhatov, 2000, 1991).

Creation of the Kazakh fat tail semi-coarse breed was begun by V.A. Balmont using complex reproductive breeding of local Kazakh coarse-haired sheep with Edilbay, Saraja, Degeress and Tadzhik breeds (Balmont, 1968).

Here we present the results of a pilot research project for the introduction of massive SNP genotyping of sheep raised in Kazakhstan. We applied Illumina Beadchip technology for the first time to perform SNP genotyping of local Kazakh sheep breeds. We investigated the genetic diversity and population structure of five local Kazakh sheep breeds (Edilbay, Akzhayik, Saryarka, Kazakh fine-wool, Kazakh semi-coarse wool) using the Illumina OvineSNP50 panel. Furthermore, we compared these data with the Sheep HapMap project dataset compiled by the International Sheep Genome Consortium to compare variability in populations of local breeds in Kazakhstan with global domestic sheep diversity. This work will lay a foundation for the introduction of genomic methods to sheep husbandry in Kazakhstan and can be a basis that can be extended to other branches of animal and plant agriculture in Kazakhstan.

## Materials and methods

### Collection of samples and DNA extraction

Samples from Edilbay sheep were collected on the “Birlik” (n=794) and “Azhar” (n=470) farms in West Kazakhstan. Samples from Akzhayik sheep were collected on the “Atameken” (n=1110), “Kuanysh” farm (n=500), and “Saltanat” farms (n=410), also in West Kazakhstan. Samples from Kazakh fine wool breed were collected in the Almaty region (624 samples -”PZ” R-Kurty”). For Kazakh fat-tailed semi-coarse-wool sheep, samples were taken on the Altyn Asel farm (n=270) in the Aktobe region, the Otkanzhar farm (n=531) in the Karaganda region and the Khasiev (n=452) and KaraAdyr farms (n=502) in East Kazakhstan. Samples for Saryarka sheep were collected on the Sarysu farm (n=498) in the Karaganda region.

Plucks from sheep ears were collected for DNA extraction in Eppendorf tubes, fixed with ethanol and stored at −20 °C. Before processing, the samples were washed twice with phosphate buffered saline to remove fixatives. DNA was extracted using MasterPure Complete DNA & RNA Purification Kits (Lucigen Simplifying Genomics, USA) according to the manufacturer’s protocol and diluted to 50 ng/μL.

### SNP genotyping and quality control

SNP genotyping was conducted using an Illumina iScan System with ovine SNP50 BeadChips according to the manufacturer’s protocol. HiScan raw intensity data (IDAT) were loaded into the Genotyping module of GenomeStudio v.2.0.4 software (Illumina, 2016) for genotype calling and quality conrol. Genotypes were inferred using manifest and cluster definition files accessed from the manufacturer’s website (accession date 08.07.2019). After initial SNP clustering and statistical evaluation, samples having a median GC score and call rate of < 0.9 were excluded, and the SNP statistics was recalculated. All data that remained after filtering (i.e., 50% GC ≥ 0.9 and call rate ≥ 0.9 for samples and SNP) were exported in a PLINK input format (ped+map) using the PLINK Input Report Plug-in v2.1.4 for GenomeStudio. SNP having minor allele frequency (MAF) < 0.02 and deviations from Hardy-Weinberg equilibrium at a critical *p*-value level of 0.0001 were filtered out by PLINK 1.9 (Purcell and Chang, 2015).

### Data analysis

General population statistics including expected and observed heterozigosity, Write’s fixation index (Fst) and linkage disequilibrium test were calculated by with PLINK1.9 and then processed with R (R Core Team, 2019). Principal component analysis (PCA) was performed using the adegenet package in R (Jombart, 2008; Jombart and Ahmed, 2011). Model-based analysis of population structure was conducted using two independent algorithms implemented in ADMIXTURE (Alexander et al., 2009; Alexander and Lange, 2011) and FastStructure (Raj et al., 2014) software. Both programs were run for K’s from 2 to 20 with 10 repetitions for cross validation. FastStructure was run in simple prior mode. Results were visualized with CLUMPAK (Kopelman et al., 2015). To determine the relationship of Kazakh breeds with other sheep breeds across the world, diversity data were compared with the Sheep HapMap dataset (SHM) compiled by the International Sheep Genome Consortium (Kijas et al., 2012, 2009). The SHM dataset was retrieved upon request from https://www.sheephapmap.org/ in accordance with the ISGC Terms of Access. A subset of Sheep HapMap data that included only those breeds for which there were at least 20 specimens and available SNPs in the processed Kazakh sheep dataset was selected, and then SNP physical mapping was updated according to the Illumina OvineSNP50 manifest file used for genotyping. A total of 50 randomly selected specimens from each Kazakh sheep population were merged with HapMap data using PLINK. The Ape (Analysis of Phylogenetics and evolution) package for R (Paradis et al., 2004) was used to calculate Neighbor-joining tree of distances, which was visualized further by FigTree (Rambaut, 2018). R and bash scripts describing conducted analyses are available in the GitHub repository [https://github.com/ASPozharsky/kz_ovine_SNP].

## Results

Out 6,756 samples and 54,241 SNPs, 4,660 and 30,597, respectively, satisfied the selection criteria. An additional 4,063 SNPs that had low MAF and/or deviated from HWE (p<0.0001), as well as markers having an undetermined genomic position and non-autosomal markers were excluded such that 26,534 SNPs were considered for further analysis. Percent of individuals passed quality criteria differs depending on population (see Table S1 in Supplementary materials). The best sample quality was from Birlik farm (Edilbay), Kara Adyr (Kazakh Semi-coarse) and Kuanysh farms (Akzahayik). The most individuals were discarded from the Otkanzhar and Altyn Asel farms (Kazakh Semi-Coarse). The low quality of these two populations indicates serious violation of storage and transport of animal material conditions by corresponding farms. However, 59 and 78 samples, respectively, were selected as suitable for further analysis.

First, general genetic diversity statistics including He, Ho, and Fst were determined for the different breeds (Table 1). All populations had similar Ho values (average 0.385), which did not substantially differ from He. The lowest within-breed Fst was between the two Edilbay populations (0.0052). Pairwise Fst across populations of Kazakh Semi-coarse wool sheep varied between 0.0095 (Altyn, Asel and Otkanzhar) and 0.0193 (Kara, Adyr and Otkanzhar). Fst between breeds did vary and allowed the population to be separated into two groups that had lower (Edilbay, Saryarka, Kazakh Semi-coarse wool) and higher (Akzhayik, Kazakh Fine-wool) inter-breed values (Fig. 1a). Separate Fst values by chromosome showed a similar distribution, except for chromosome 16 (Fig. 1b-d, Table S2 in Supplementary materials).

**Table 1.**
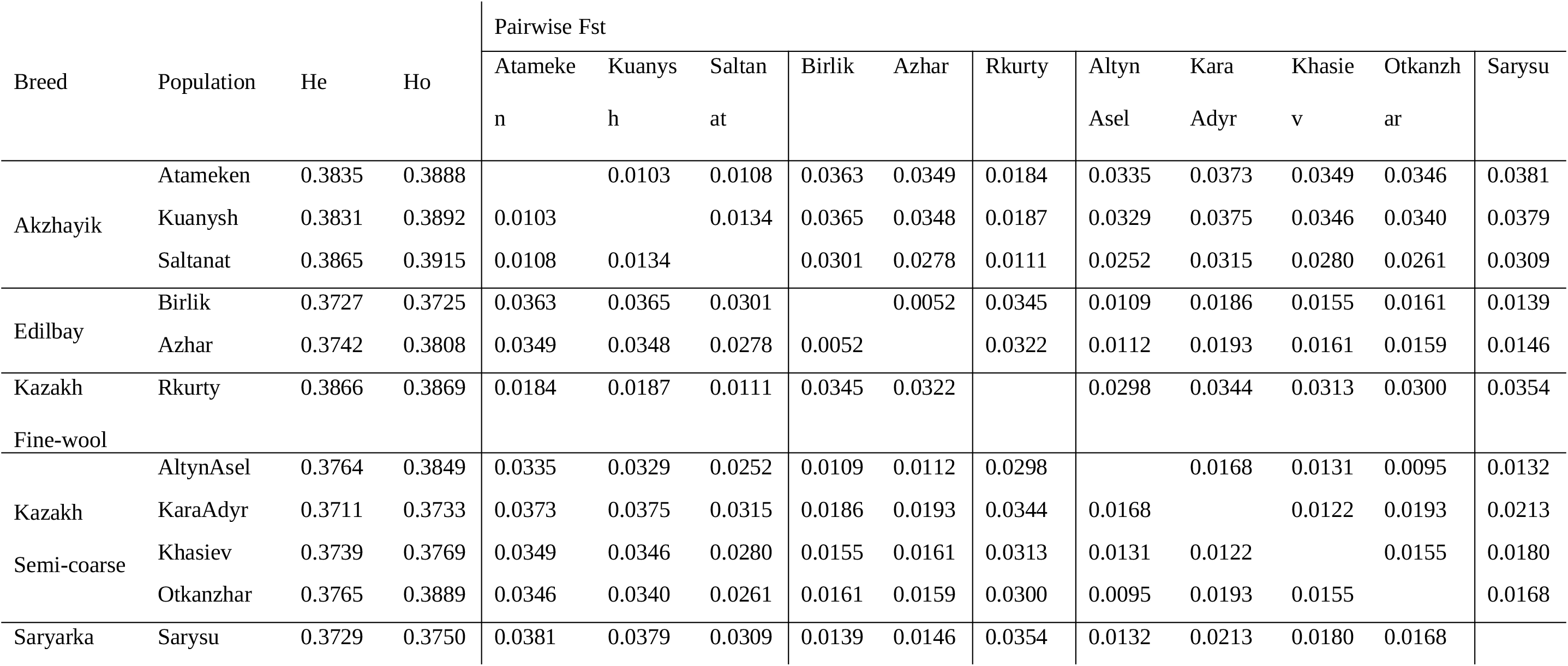
Expected and observed heterozygosity and Write’s Fst for sheep population from different farms in Kazakhstan

**Figure 1.**
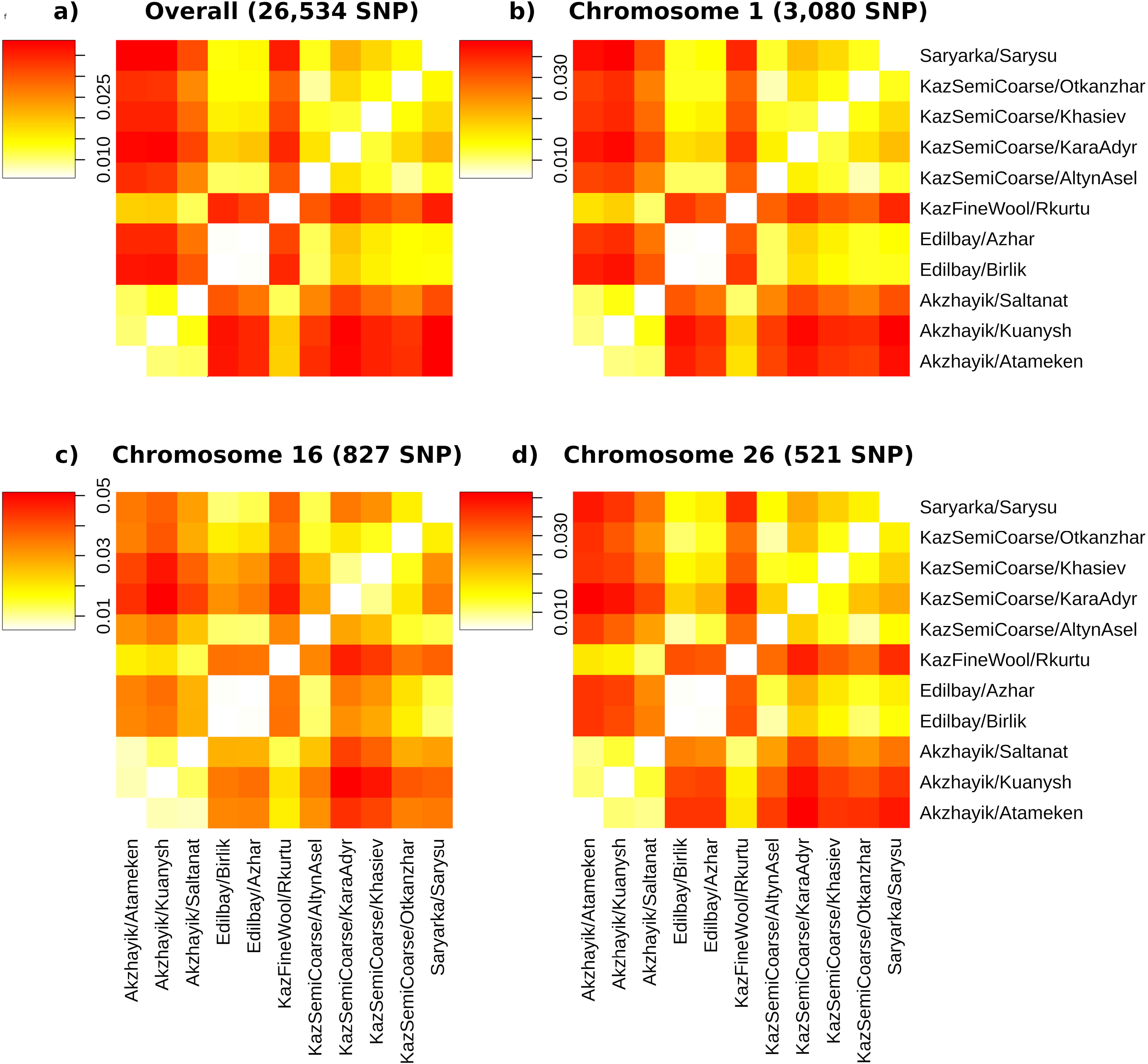
Heatmaps of pairwise Fst across Kazakh sheep populations

Linkage disequilibrium decay was evaluated separately for each breed and each population and expressed as an average R2 coefficient for 20 kb intervals (Fig. 2). Akzhayik, Edilbay and Kazakh Semi-coarse wool sheep had similar LD decay rates that were close to that for the entire sample. For Edilbay, Akzhayik and Semi-coarse wool breeds, LD decay can be compared between populations. Two Edilbay populations had very similar decay rates. Among Akzhayik sheep, the Saltanat population showed different LD decay from Kuanysh and Atameken. All populations of the semi-coarse wool breed had different LD decay rates.

**Figure 2.**
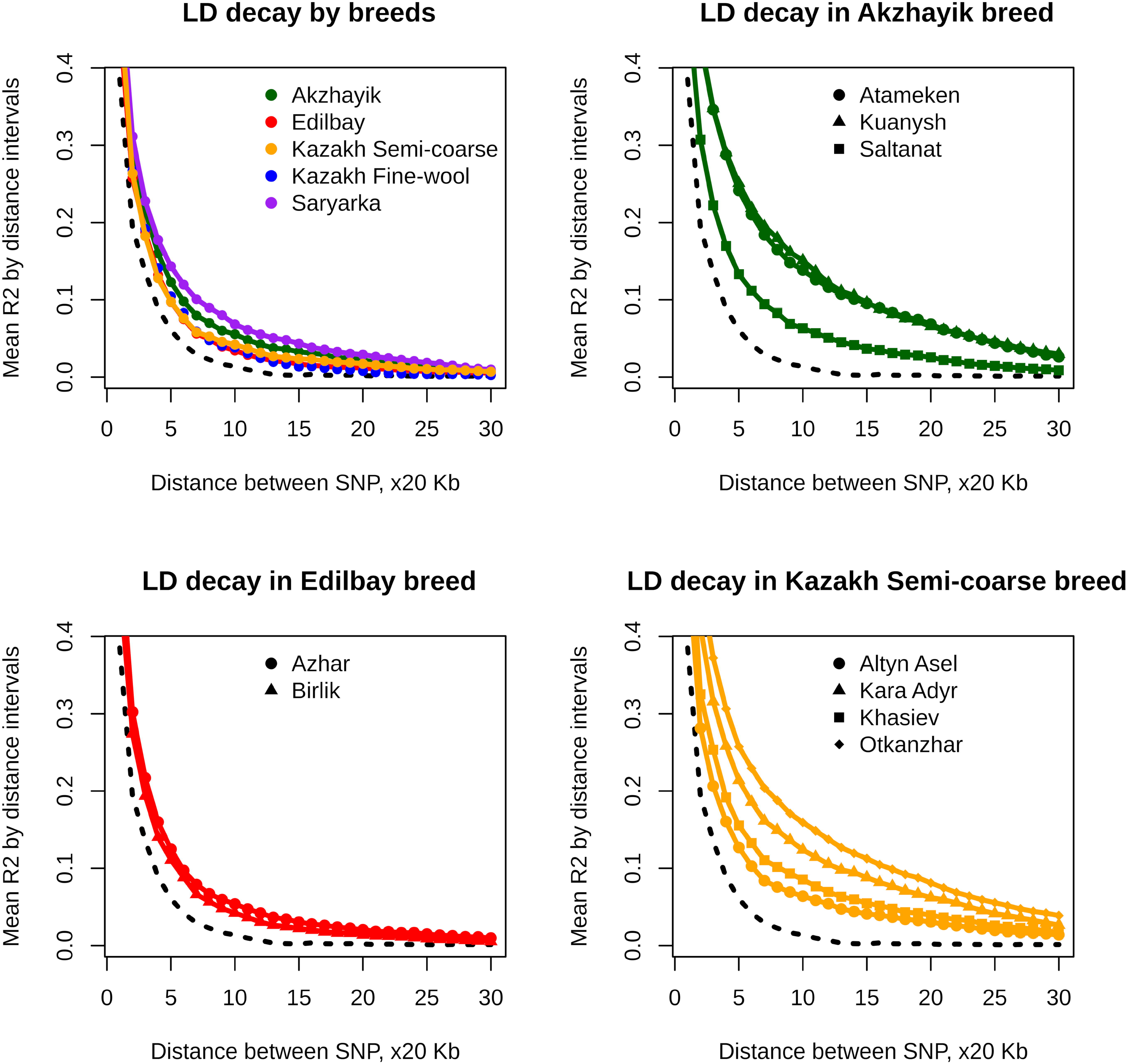
Linkage disequilibrium decay between breeds and populations of the same breed (Akzhayik, Edilbay, Kazakh Semi-coarse breed). The black dashed line represents LD decay for the entire dataset.

Principal component analysis was next conducted and the first 100 components were retained. All components explained a relatively small part of the total variance (Table S3; Fig. 3B; Fig. S1). The first three PCs explained 15%, 4% and 3.6% of variance, respectively (Fig. 3c-e). The first PC revealed two major clusters that corresponded to the abovementioned groups formed according to Fst. Several components had smaller differences between breeds and populations within these two groups (Fig. 3a).

**Figure 3.**
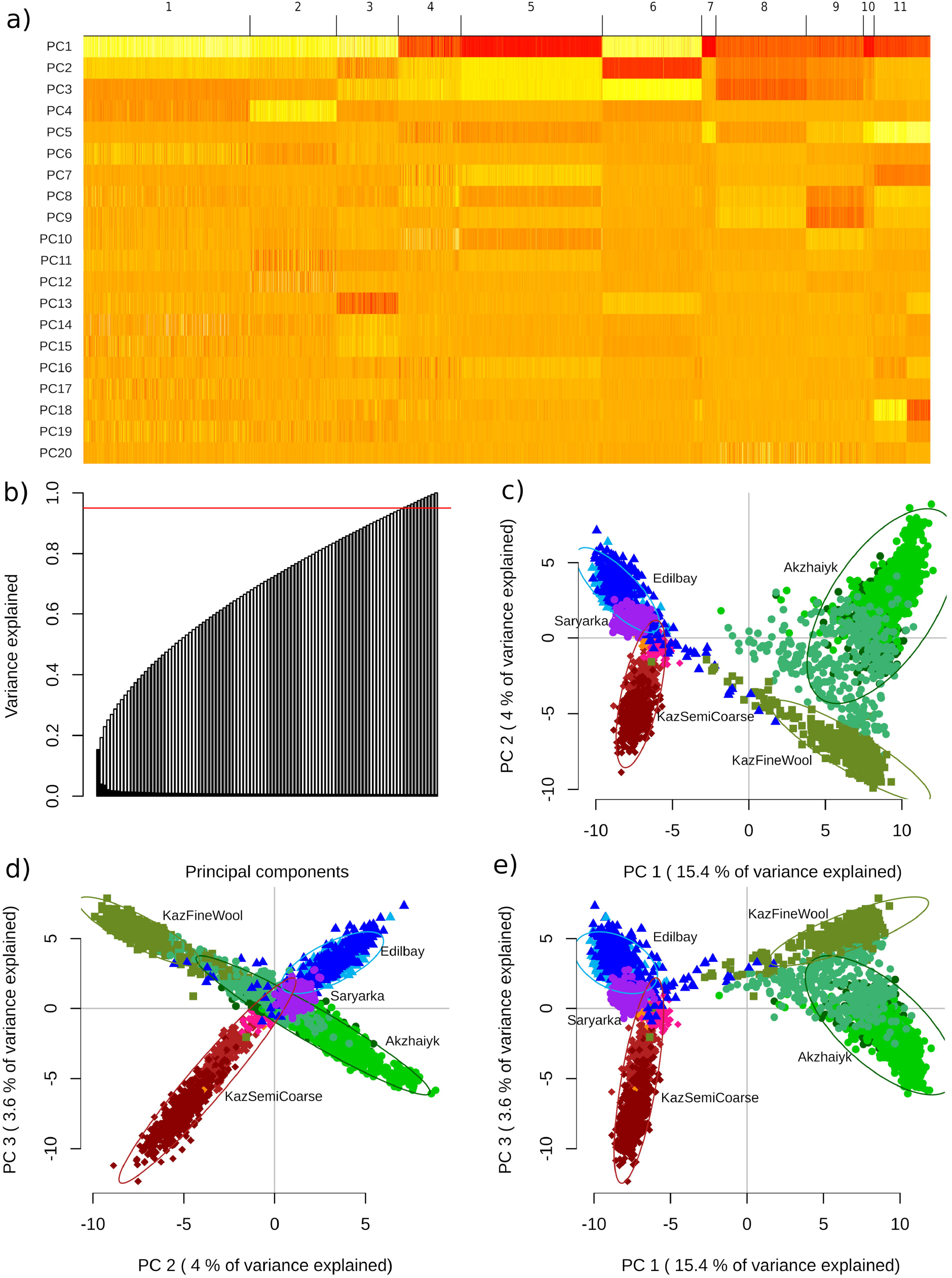
Principal Component Analysis of Kazakh sheep populations. a) Heatmap of first 15 components. Populations are designated as follows: Akzhayik: Atameken (1), Kuanysh (2), Saltanat (3); Edilbay: Azhar (4), Birlik (5); Kazakh Fine-wool sheep – R-Rurty (6); Kazakh Semi-coarse wool sheep: Altyn Asel (7), Kara Adyr (8), Khasiev (9), Otkanzhar (10); Saryarka – Sarysu (11). b) Plot of cumulative variance explained by PCs. 95% level is shown by the red line. c,d,e) Pairwise 2D dot plots of the first three components.

ADMIXTURE and FastStructure were run to determine the genetic structure within and between populations and breeds. The cross validation error score did not converge to its minimal value in all runs of K values up to 20 for both algorithms. Thus, the following Ks were selected for examination: 2 (minimum), 5 (number of Kazakh breeds), 11 (number of Kazakh breed populations) and 20 (maximum) (Fig. 4; see also Figures S2 and S3 in Supplementary materials for all original plots). When K=2, both algorithms produced essentially identical results. As with diversity features, these calculations allowed all breeds to be differentiated into two clusters: one having Edilbay, Saryarka and Kazakh Semi-coarse wool sheep, and the second having Akzhayik and Kazakh Fine-wool sheep. Additional K results differed between the two programs. At K=5, FastStructure results showed more mixed populations than ADMIXTURE and found effectively no within-population genetic structure with further models. The general population structure revealed by ADMIXTURE was more stable.

**Figure 4.**
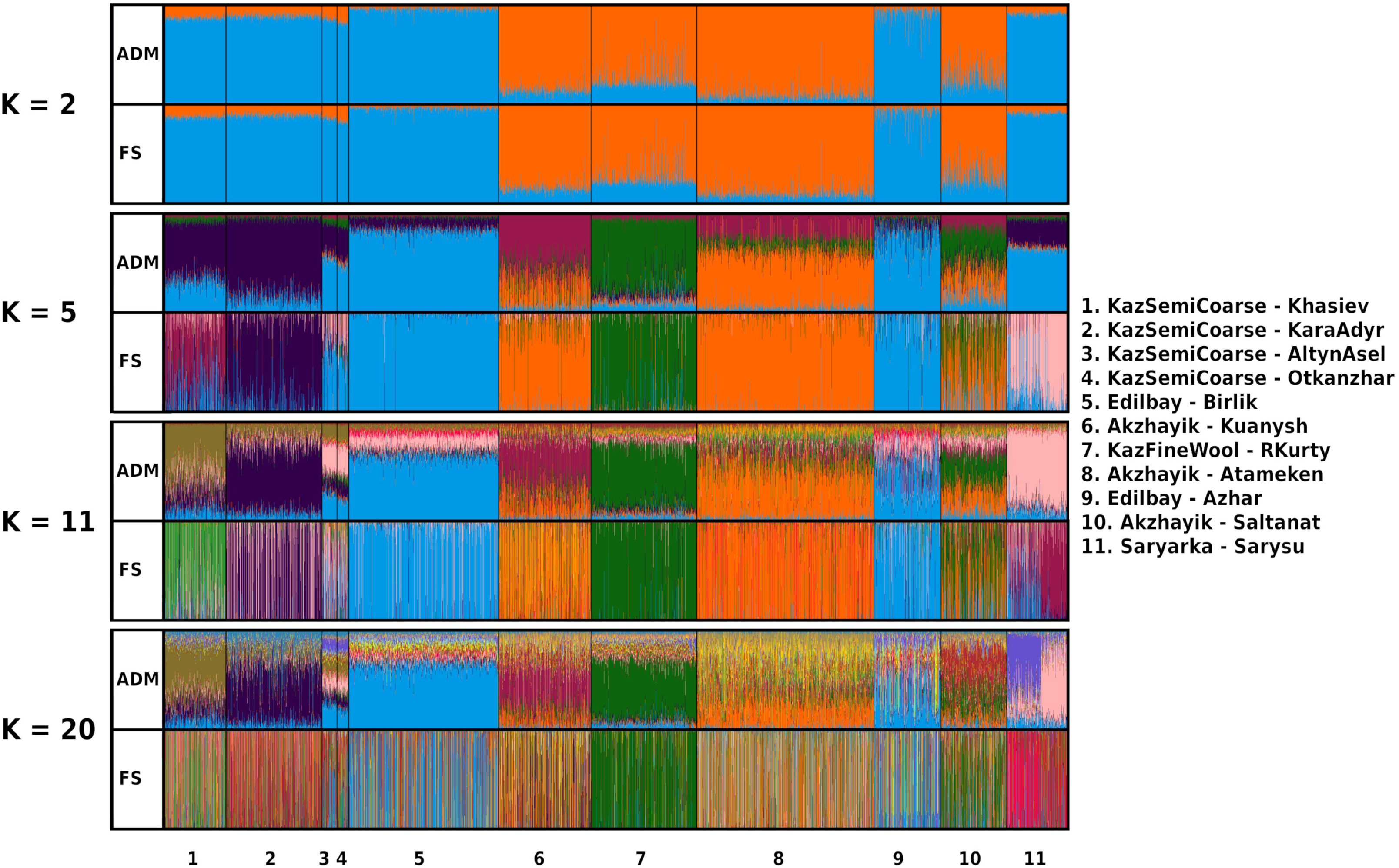
Model based population structure analysis using ADMIXTURE (ADM) and FastStructure (FS) software.

Populations of sheep that were the same breed but came from different farms showed high heterogeneity both within and between populations. Akzhayik and Kazakh Semi-coarse breeds had a highly mixed gene pool in every sampled population. Meanwhile, the Edilbay population from Azhar was more mixed compared to the Birlik population.

To compare diversity of Kazakh sheep breeds to breeds worldwide, we built a genetic distance tree using the Neighbor-joining method. All Kazakh breeds are shown as a distinct cluster within the entire tree (Fig. 5). Edilbay is a heterogenous basic group, whereas other breeds appear subsequently as separate branches. The breed clusters diverged from Edilbay in the following order: Saryarka and Kazakh Semi-coarse sheep, Akzhayik and Kazakh Fine-wool sheep, followed by additional Asian and South-West Asian breeds.

**Figure 5.**
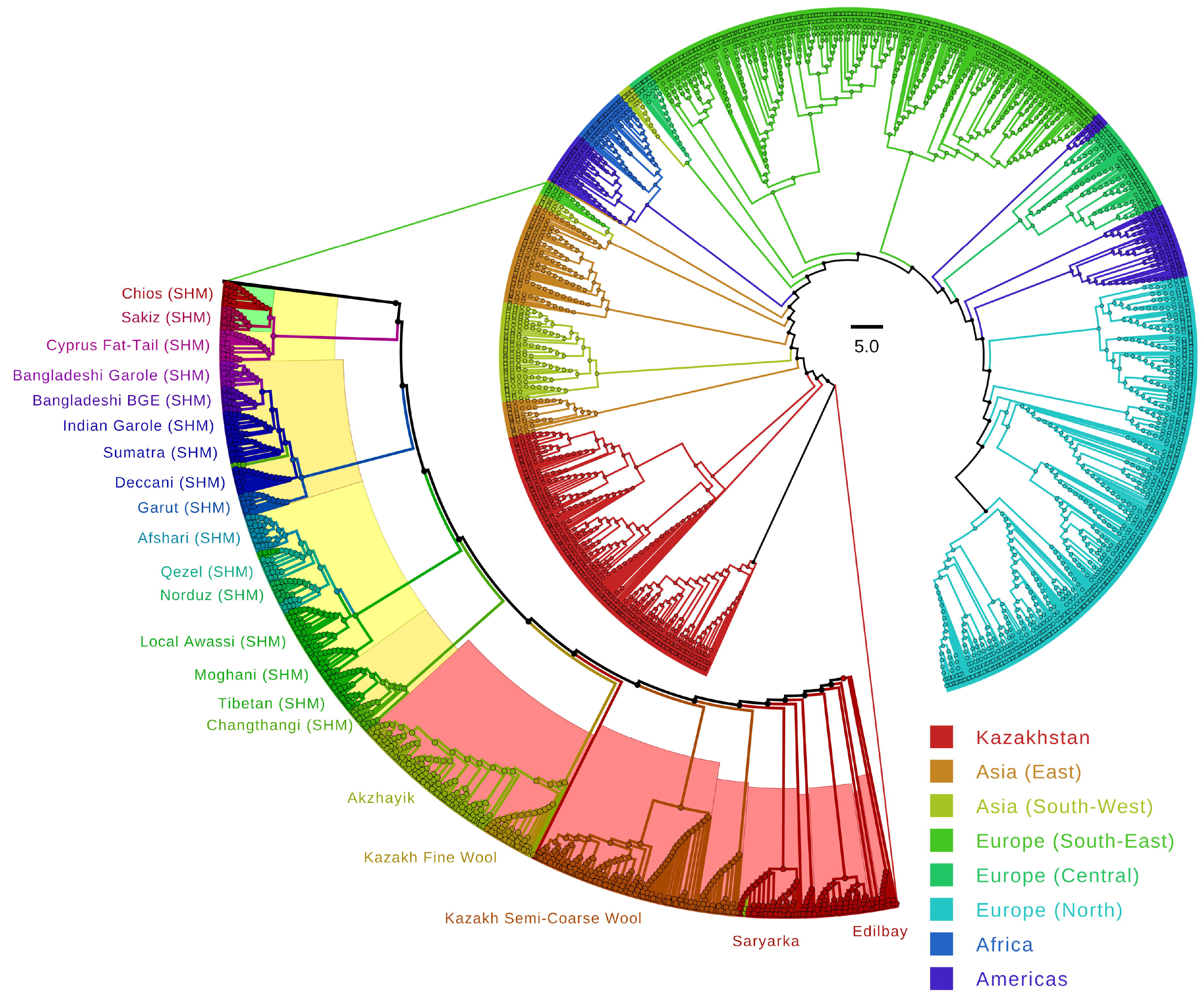
Neighbor-joining tree of genetic distances across Kazakh sheep breeds and breeds in the Sheep HapMap (SHM) dataset. All geographical annotations of SHM breeds are in accordance with ISGC sample annotations.

## Discussion

Comparison of genetic structures between the studied breeds was consistent with the known history of the respective breeding processes. Edilbay and its descendants, Saryarka and Kazakh Semi-coarse wool sheep, were shown to be distinct from Akzhayik and Kazakh Fine-wool sheep. On the other hand, variations in breeds and populations within these groups were less clear. PCA results demonstrated that the most discriminating first component could explain only 15% of the total variation, whereas other components described differences between breeds and population even less well. All remaining components tended asymptotically to the level of 0.5% of explained variance. However, these values are relatively high with respect to world-wide diversity data (Kijas et al., 2009), which are in agreement with the weak population structure reported for mitochondrial and genomic SNP data (Kijas et al., 2012; Meadows et al., 2005). As the ADMIXTURE and FastStructure analysis results showed, all populations in this study tended to separate from each other. However, neither method were not able to reached an optimal configuration with the lowest cross-validation error in runs having K=20. Although FastStructure was shown more efficient in resolving population structure (Raj et al., 2014), with this method noise increases rapidly with increasing K. Thus, FastStructure appears to be too sensitive to variation, with individual variability overwhelming population integrity. Meanwhile, the ADMIXTURE results were useful for further observations and conclusions, although in the presence of population variations the genetic structure was less clear with increasing individual values for K. The Kazakh Fine-Wool sheep population and Birlik population of Edilbay sheep retained high homogeneity even at maximal K. Akzhayik and Semi-coarse breeds had high heterogeneity between populations from different farms, which was also supported by varying LD decay rates. One noticeable observation for the Semi-coarse breed populations is that the subset of SNPs on chromosome 16 had a differing pattern for between-population Fst values (Fig. 1). Populations from Khasiev and Kara Adyr farms differed more not only from other breeds, but also from the Otkanzhar and Altyn Asel populations of the same breed. This finding might imply that all or a significant part of chromosome 16 was introduced from the genetic pool during the breeding process, and lies outside of relationships with other studied breeds. The low number of SNPs included in this study did not allow us to determine the exact origin of chromosome 16, so additional investigations are needed to clarify this finding. Edilbay populations from Birlik and Azhar farms had the lowest Fst value across all population pairs and practically identical LD decay rates. The Birlik population was the most homogeneous group, which was supported by PCA. The Azhar population is more mixed, but still expressed high similarity to Birlik. We had only one population for both Saryarka and Kazakh Fine-wool breeds and thus could draw no conclusions about within-breed populational variability. However, when high Ks (near 20) were tested, the Saryarka could be divided into two sub-clusters. This separation was also visible in PCA results, starting from PC 13, and with particular impact of PC 17 (Fig. 3a, S3). Each component in this range was responsible for no more than 1% of all variation, so ADMIXTURE demonstrates high sensitivity even to small variability components.

The neighbor-joining tree build with the addition of SHM project data allowed data for Kazakh sheep varieties to be placed in the context of global variations in sheep breeds. Considering the hypothesis that domesticated sheep have an Indo-Iranian origin, distribution of all sheep breeds should follow the historical sequence from origins in Asia with expansion into the Mediterranean region, Europe and then Africa. In such a context, the basic position of Edilbay sheep implies that this traditional lineage conserves the genetic pool of hypothetical ancestral domestic sheep varieties. This possibility is supported by previous studies on mitochondrial genetics (Hiendleder et al., 1998). However, in general there is a lack of detailed comparative genetic studies on Kazakh sheep breeds and Edilbay particularly. On the one hand, studies including Kazakh sheep varieties usually limited by a local context (Blackburn et al., 2011; Dossybayev et al., 2018; Mukhametzharova et al., 2018; Ozerov et al., 2008a, 2008b). On the other hand, in global between breed diversity studies Kazakh breeds are not being included (Kijas et al., 2012; Meadows et al., 2007, 2005) or have low representation (Hiendleder et al., 2002; Meadows et al., 2011) in global between-breed diversity studies. As such, further in-depth investigation of the importance of Kazakh sheep varieties, particularly for the Edilbay breed, in global sheep diversity is important for understanding the history of sheep domestication. Additional quantitative work involving a larger set of genome-wide markers and qualitative studies such as those that determine haplotypes using reliable mitochondrial and Y-chromosomal markers is also needed. Results from such studies will have practical importance for exploration of Kazakh sheep breeds as a previously understudied source of genetic material for selection, and also for understanding the genetic basis for standardization of Kazakh sheep varieties.

## Conclusion

Despite their unique proximity to the geographical and historical origin of domesticated sheep, Kazakh sheep breeds remain yet poorly studied in terms of genetic composition and its relationship to worldwide sheep diversity. Here we used medium-scale SNP genotyping of several sheep breeds in Kazakhstan to lay a basis for further molecular genetic studies. Investigation of the previously uncharacterized genetic pool of Kazakh sheep breeds can have implications not only for breeding practice, but for our fundamental understanding of sheep genetics and domestication history. Our data support that Kazakh breeds, particularly Edilbay, are direct descendants of historical domestic sheep ancestors.

The observed heterogeneity in sheep populations from different farms that were sampled in this study raises the issue of standardization among Kazakh sheep breeds. Given that genetic consistency is crucial for successful breeding and selection of sheep and other livestock, effort is needed to compile a detailed genetic description and “passport” for sheep populations in Kazakhstan.

## Ethical statement

Animal material was collected and provided by qualified personnel of the respective farms as part of routine veterinary care in accordance with local regulations. Collection of samples caused no harm to animal health.

## Acknowledgements

The study was funded by the Ministry of Agriculture of Kazakhstan as a part of a targeted funding program № 267 “Development of efficient breeding methods in branches of animal husbandry” The ovine SNP50 HapMap dataset used for the analysis described was provided by the International Sheep Genomics Consortium and obtained from www.sheephapmap.org in agreement with the ISGC Terms of Access.

## Supplementary materials

Table S1: Results of filtering of samples and SNPs by quality

Table S2: Wright’s fixation index (Fst), expected and observed heterozygosity in Kazakh sheep populations by chromosomes

Table S3: Explained variance of 100 principal components

Figure S1. Heatmap of first 50 principal components

Figure S2. CLUMPAK output of ADMIXTURE plots of Kazakh sheep populations

Figure S3. CLUMPAK output of FastStructure plots of Kazakh sheep populations

