## Supplementary figures and images for "SNP genotyping and population analysis of five indigenous Kazakh sheep breeds"

### Supplemental Figure 1

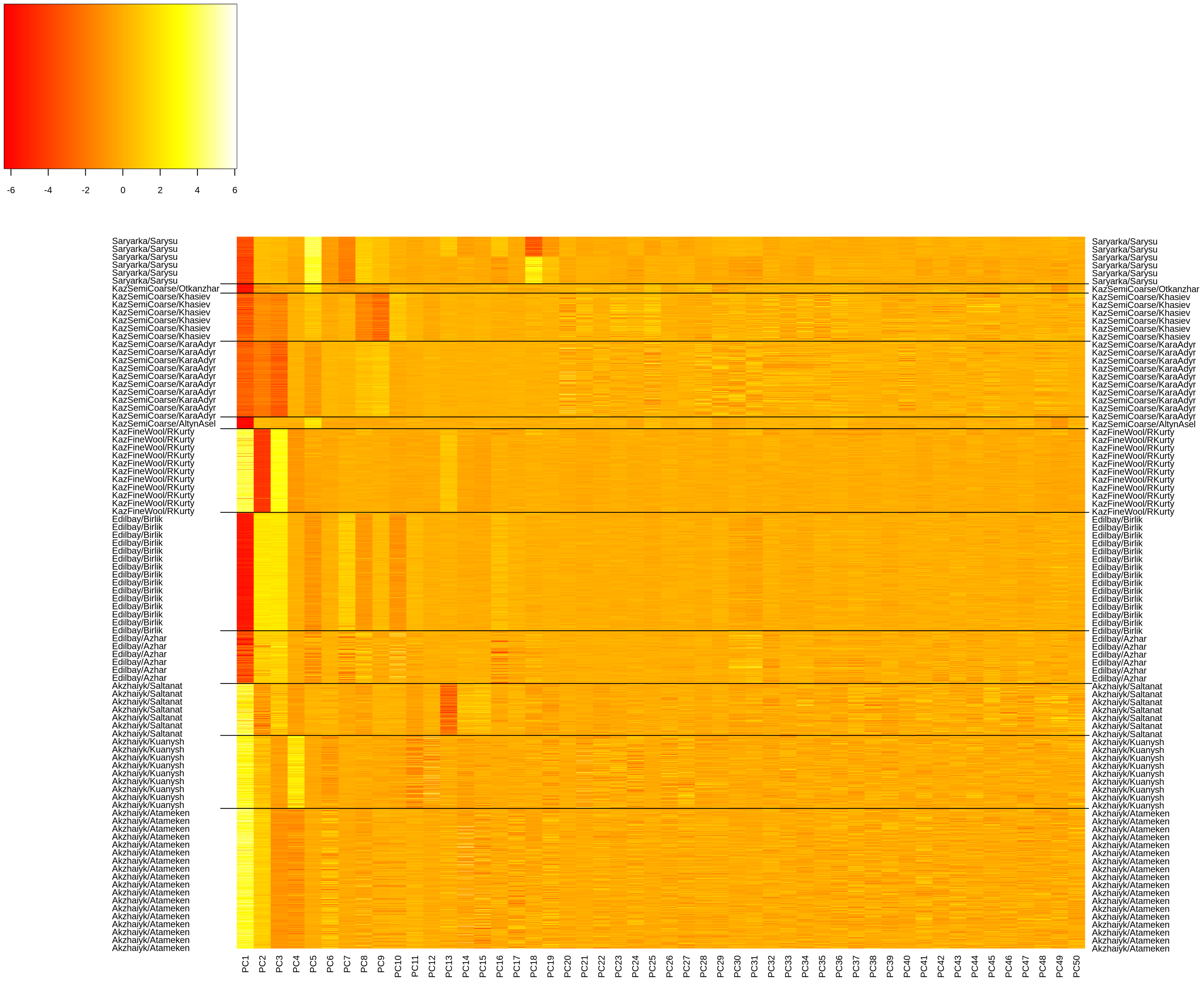
