## Supplemental Figure 2 for "SNP genotyping and population analysis of five indigenous Kazakh sheep breeds"

### CLUMPAK main pipeline - Job 1576119077 summary

#### Major modes for the uploaded data:

K=2

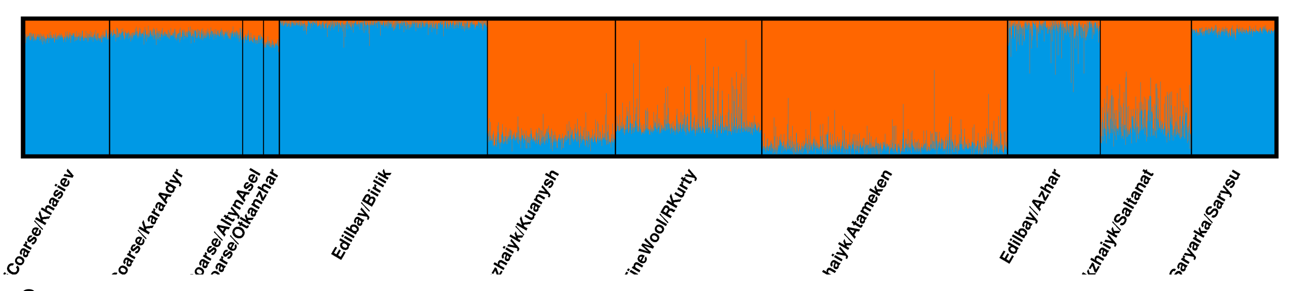

K=3

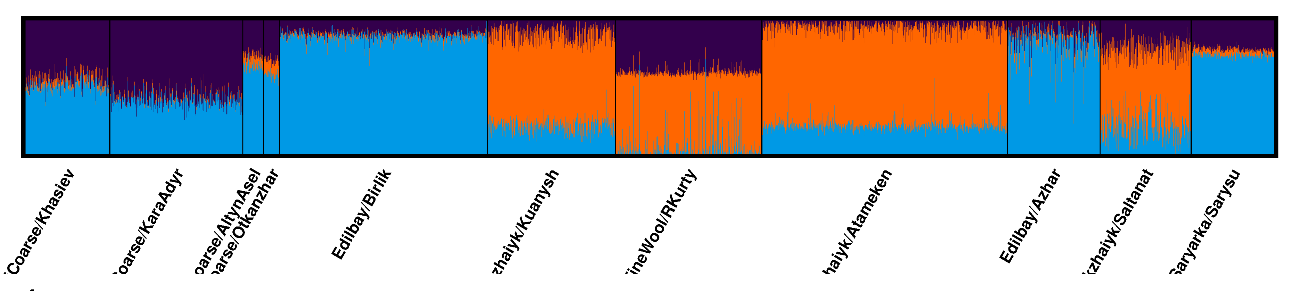

K=4

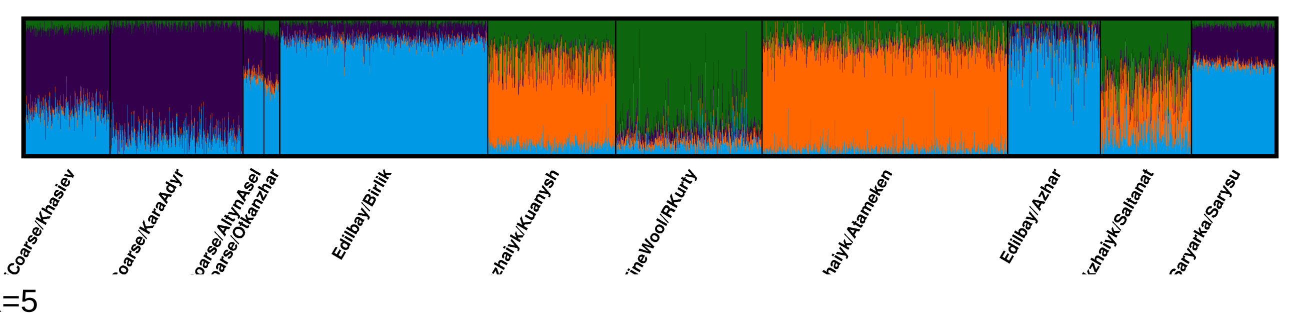

K=5

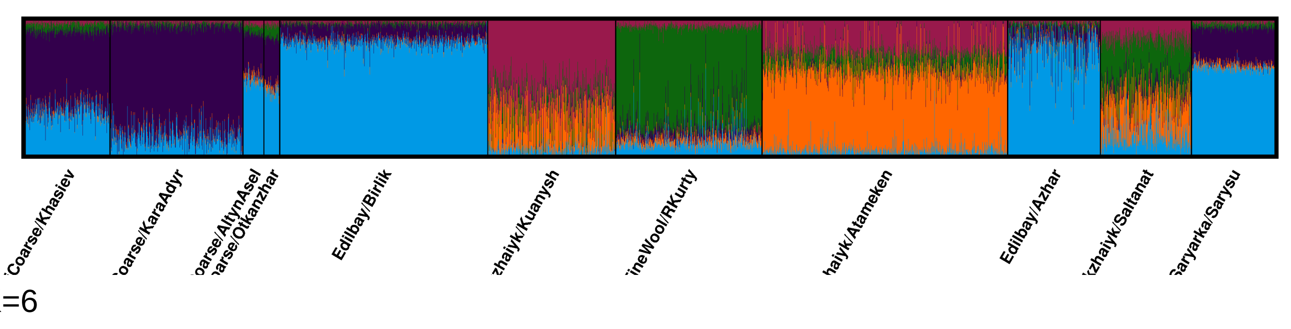

K=6

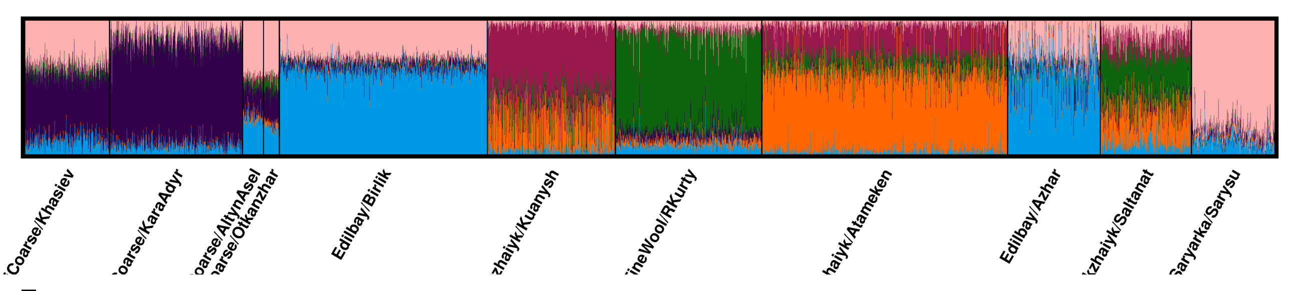

K=7

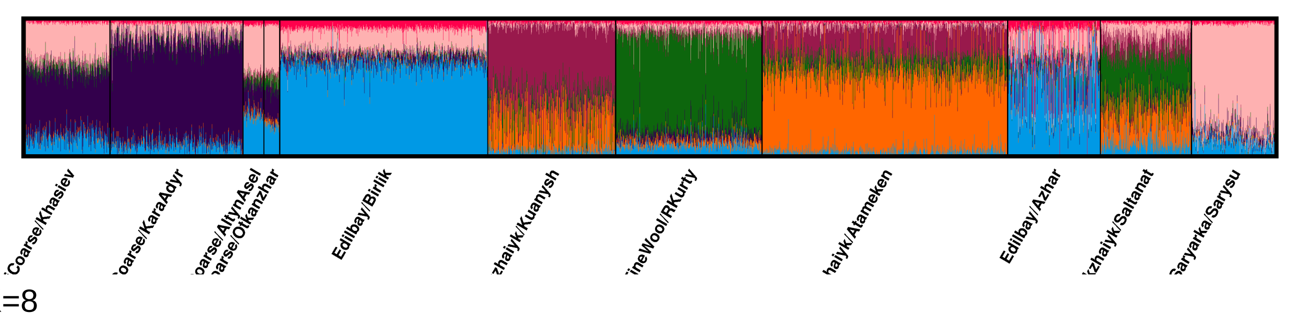

K=8

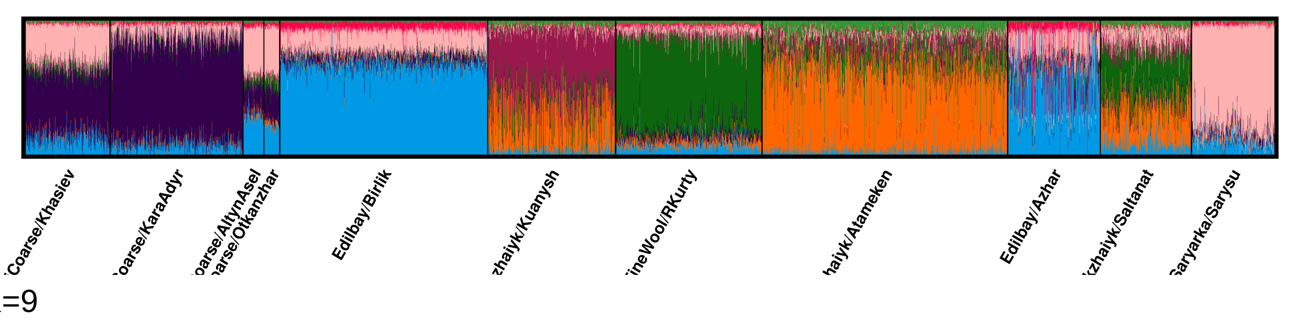

K=9

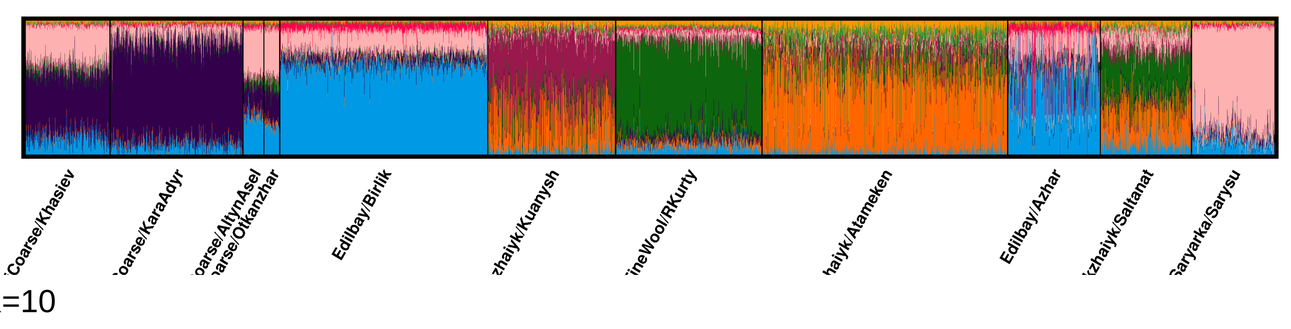

K=10

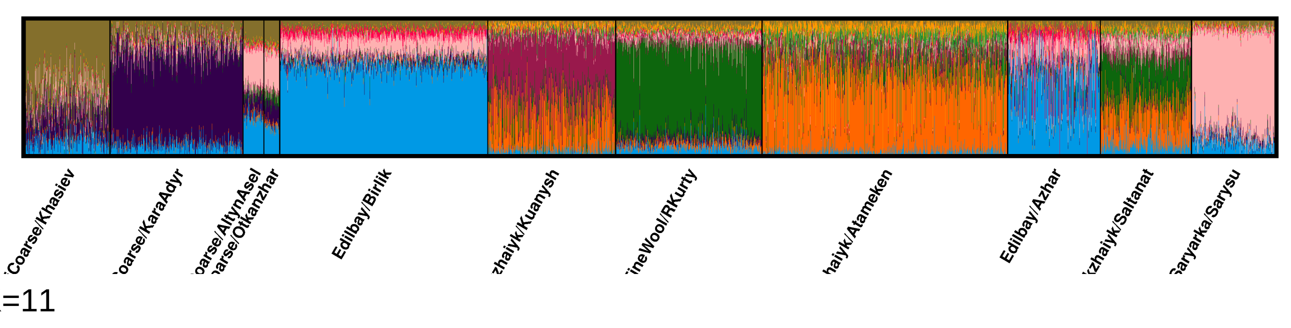

K=11

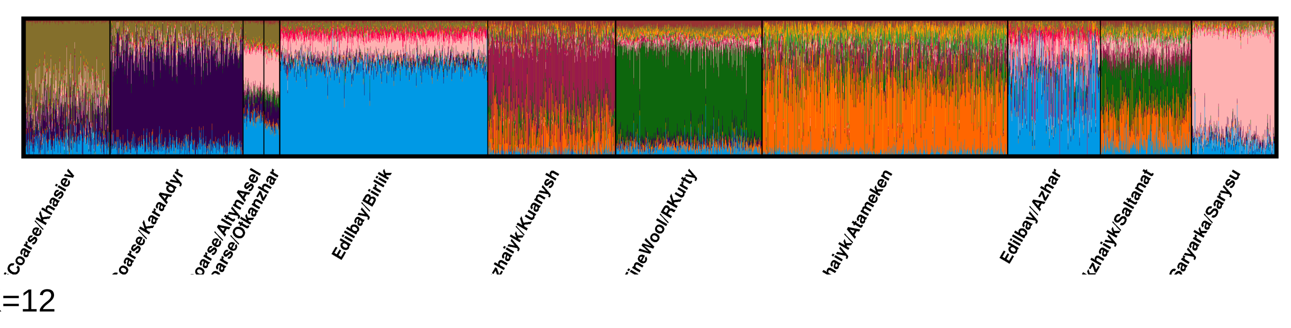

K=12

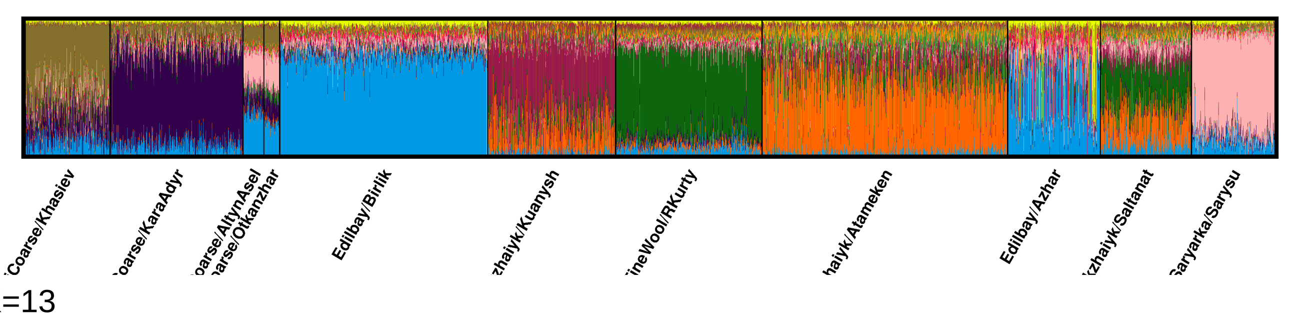

K=13

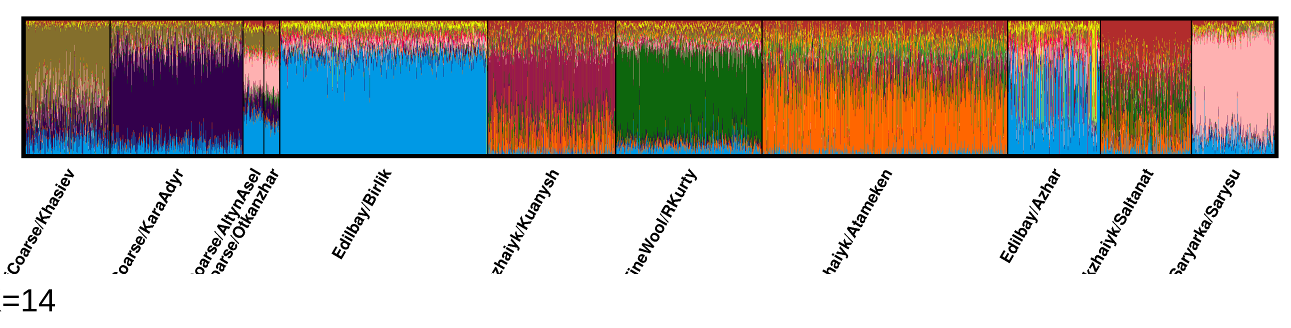

K=14

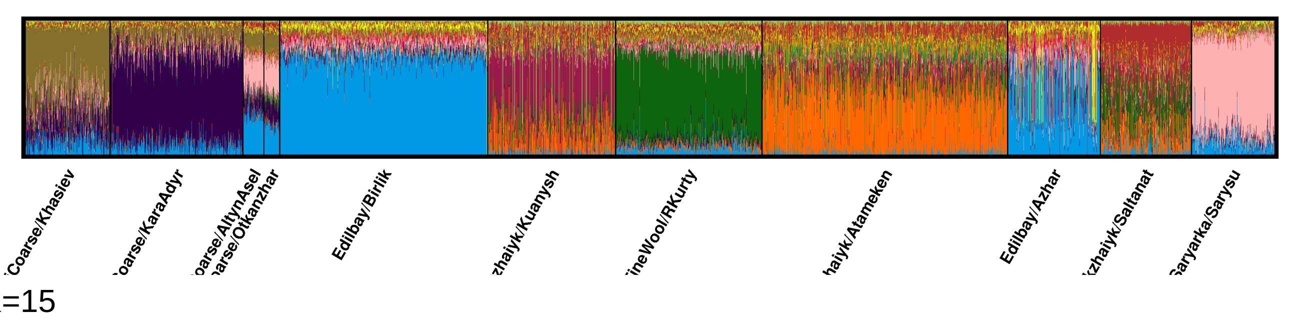

K=15

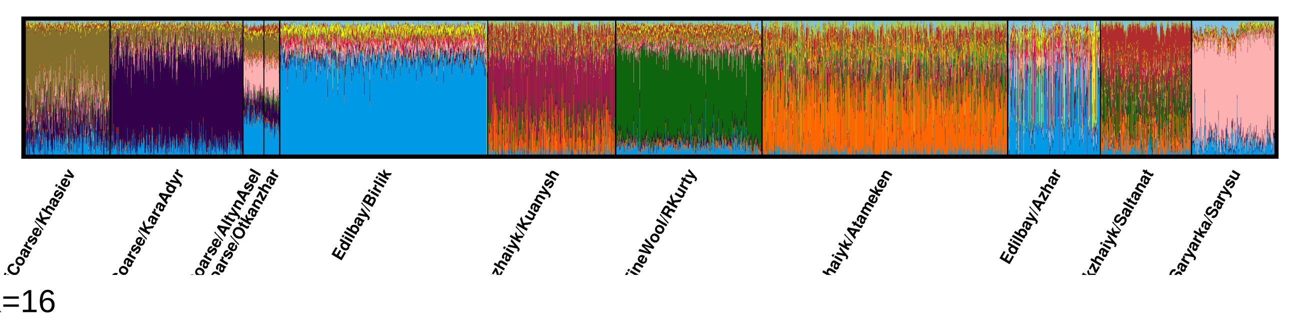

K=16

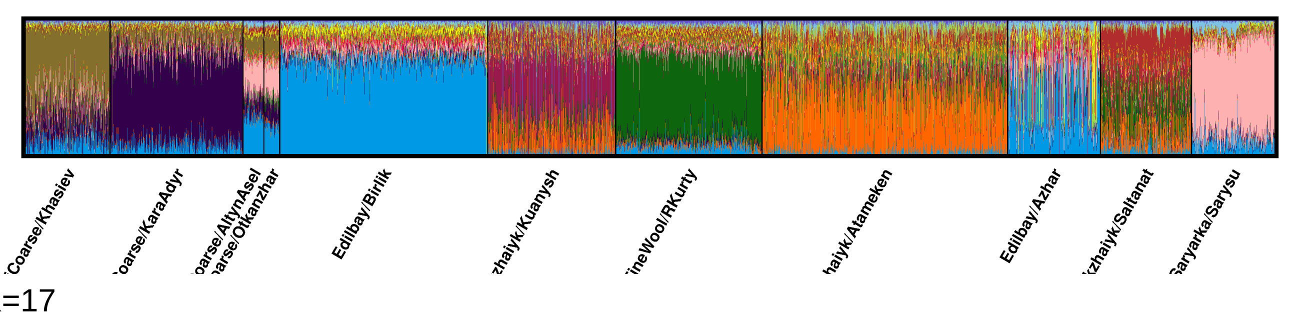

K=17

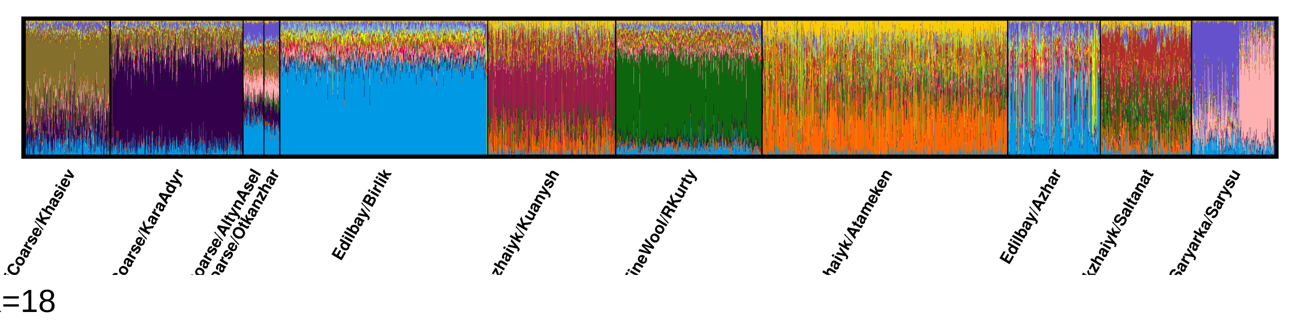

K=18

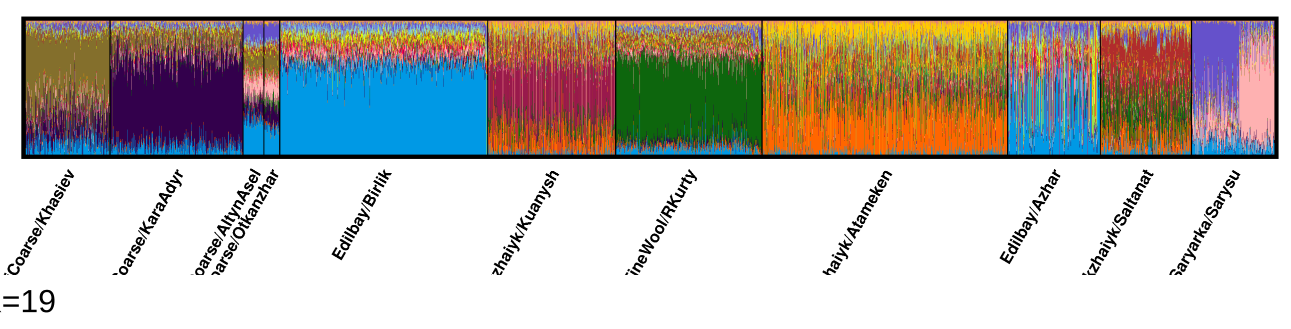

K=19

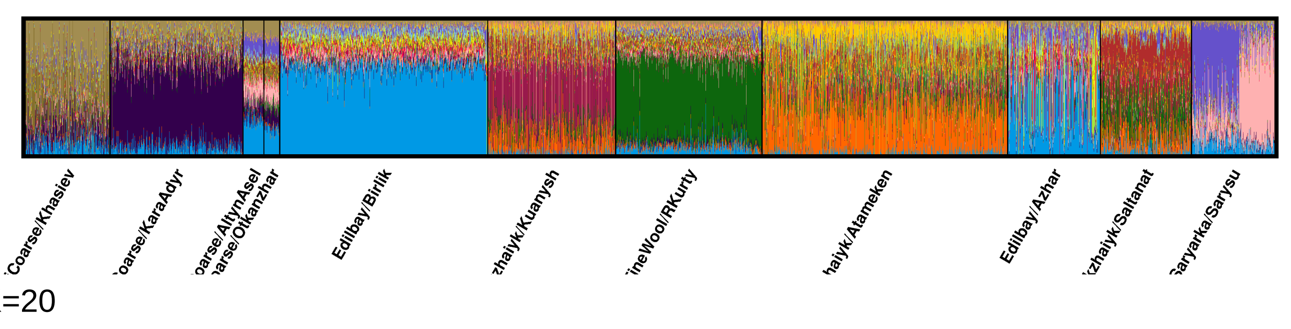

K=20

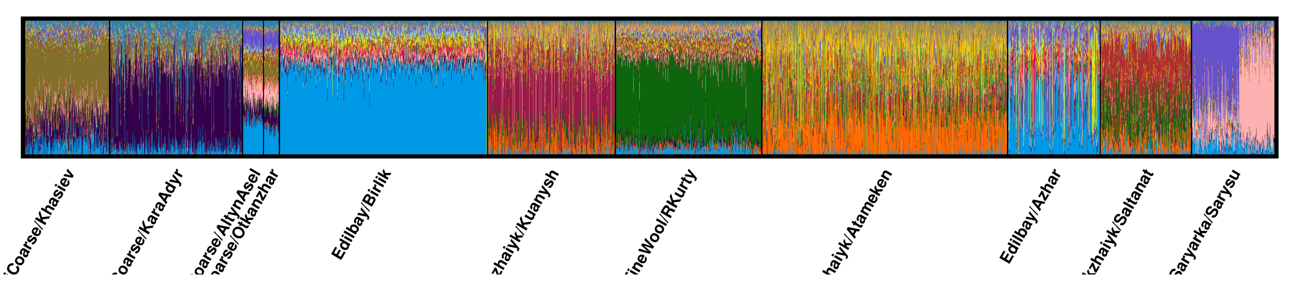

#### Minor modes for the uploaded data:

#### Division of runs by mode:

K=2 1/1
K=3 1/1
K=4 1/1
K=5 1/1
K=6 1/1
K=7 1/1
K=8 1/1
K=9 1/1
K=10 1/1
K=11 1/1
K=12 1/1
K=13 1/1
K=14 1/1
K=15 1/1
K=16 1/1
K=17 1/1
K=18 1/1
K=19 1/1
K=20 1/1
