## Supplemental Figure 3 for "SNP genotyping and population analysis of five indigenous Kazakh sheep breeds"

### CLUMPAK main pipeline - Job 1577154957 summary

Major modes for the uploaded data:

K=1

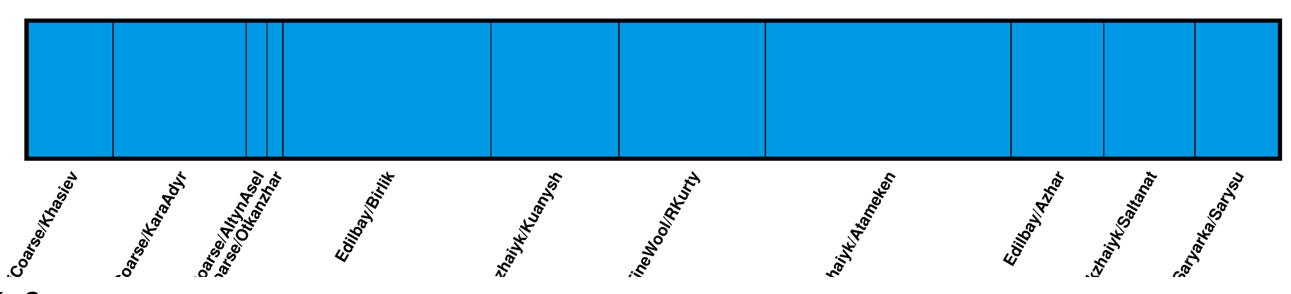

K=2

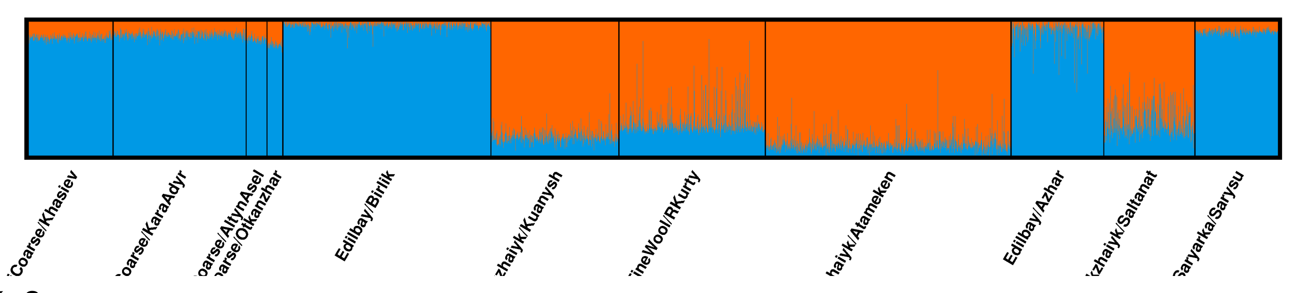

K=3

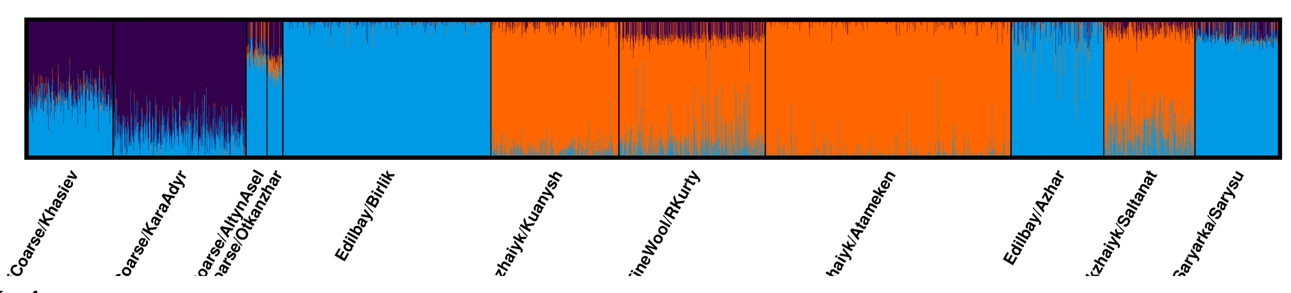

K=4

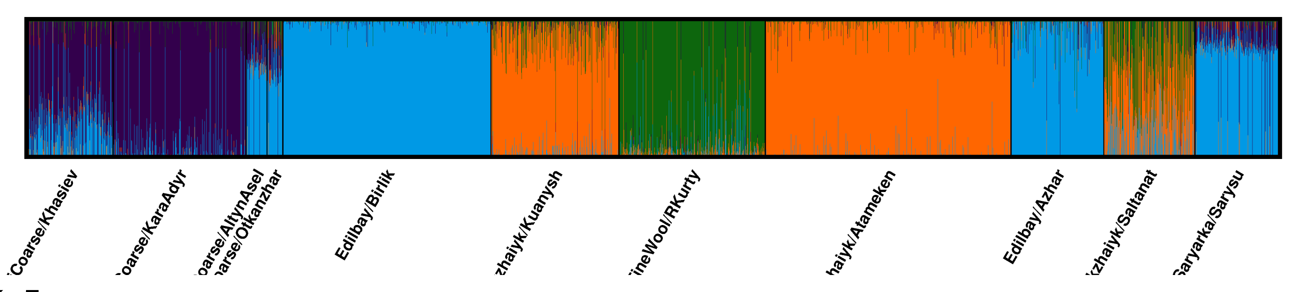

K=5

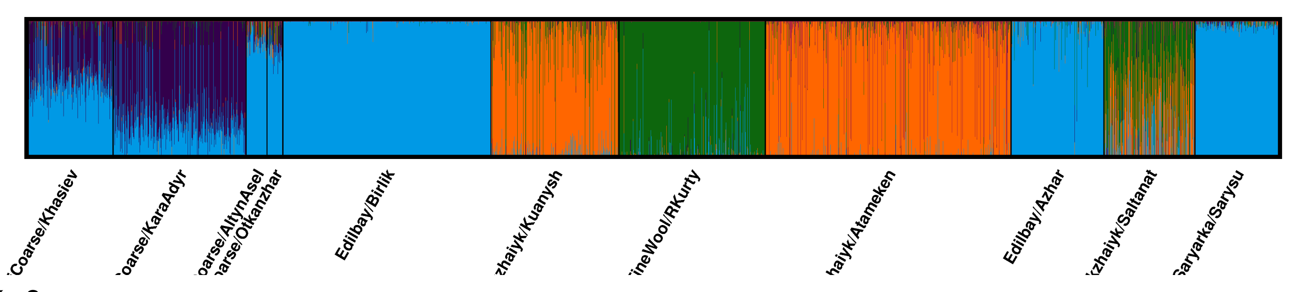

K=6

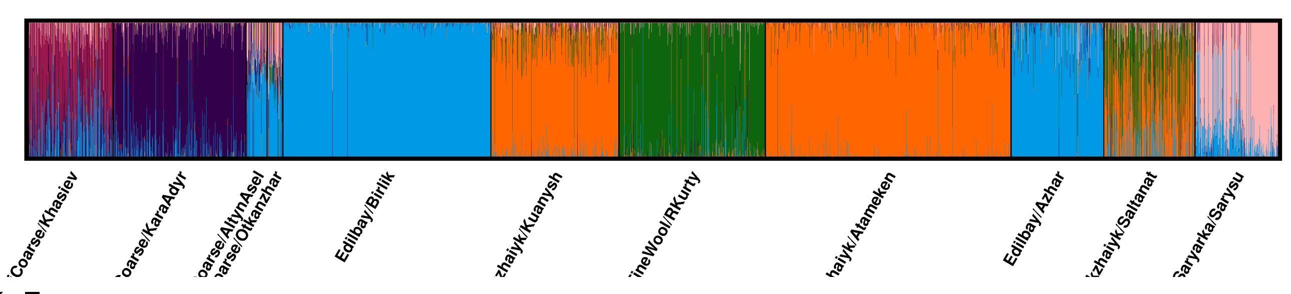

K=7

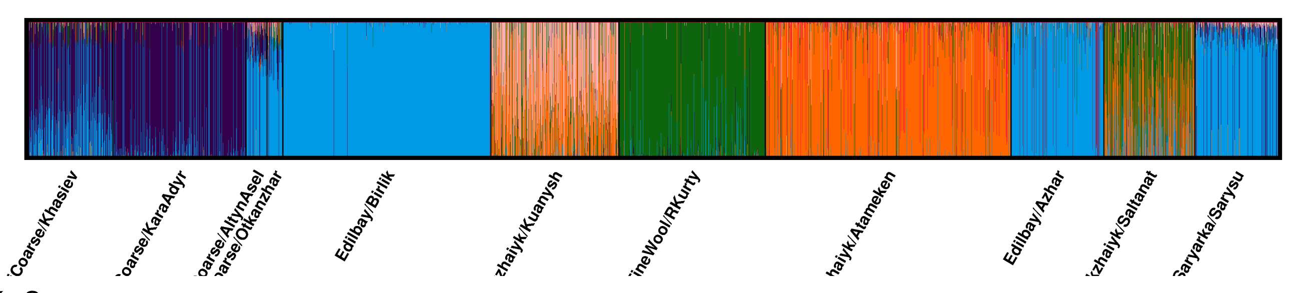

K=8

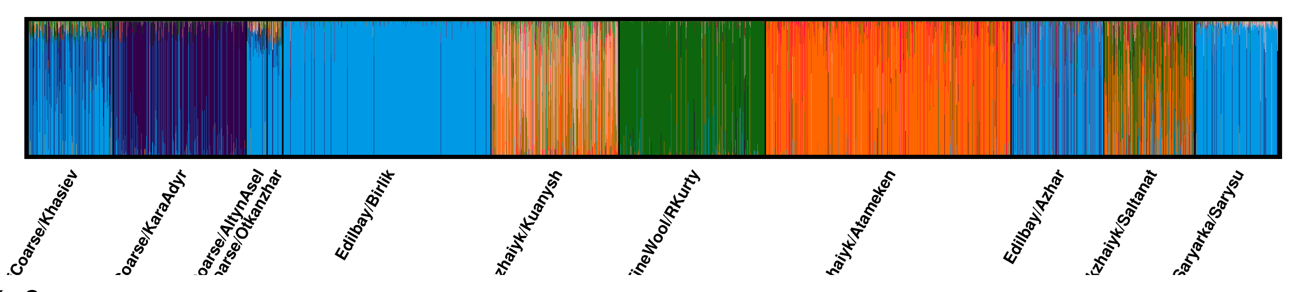

K=9

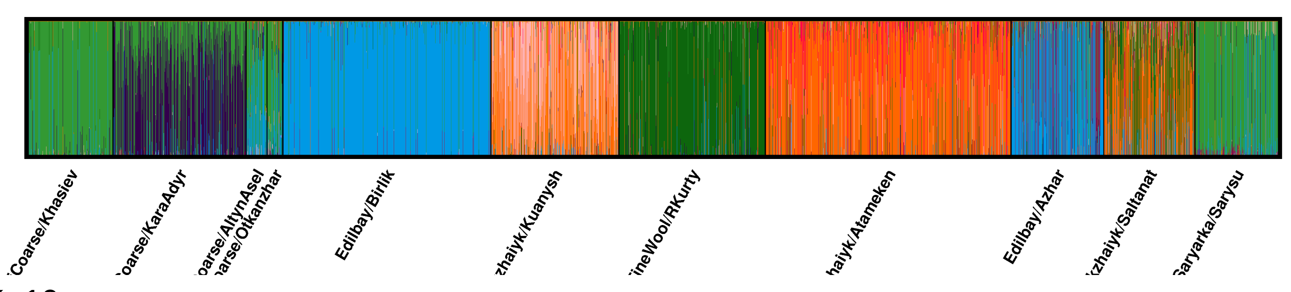

K=10

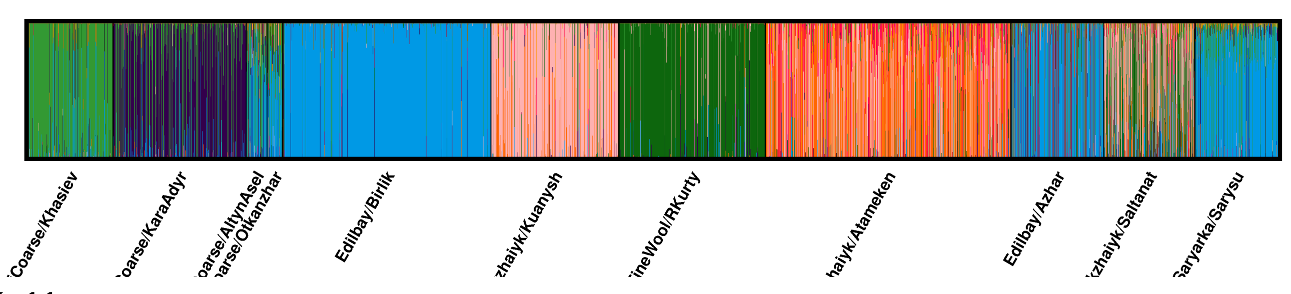

K=11

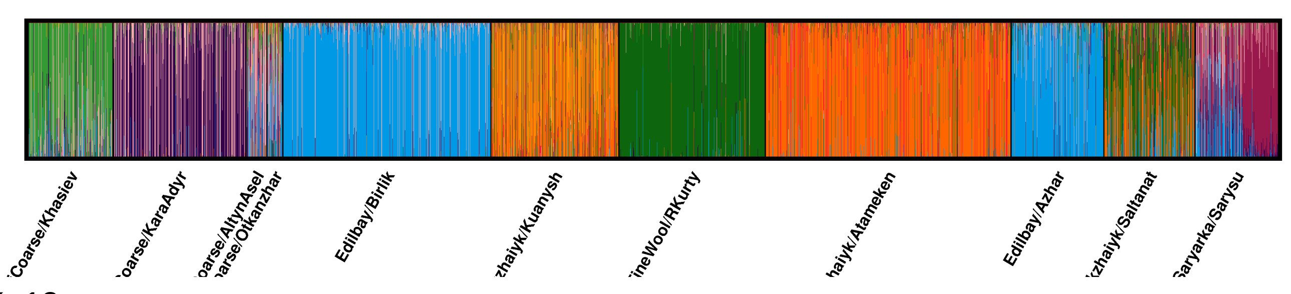

K=12

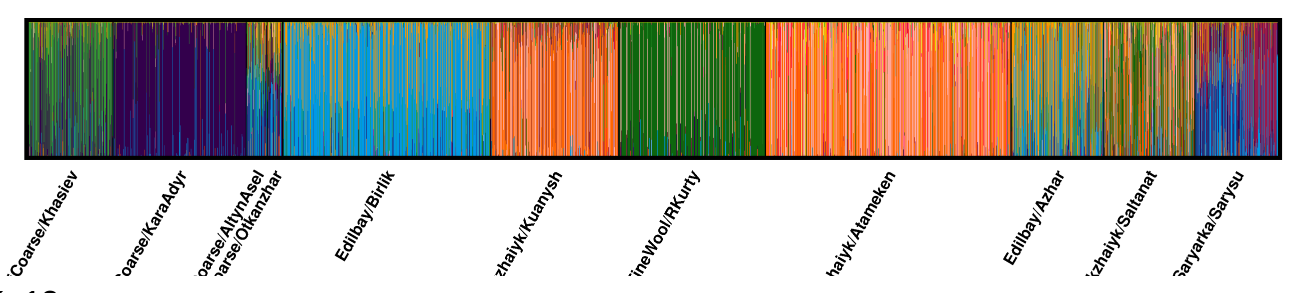

K=13

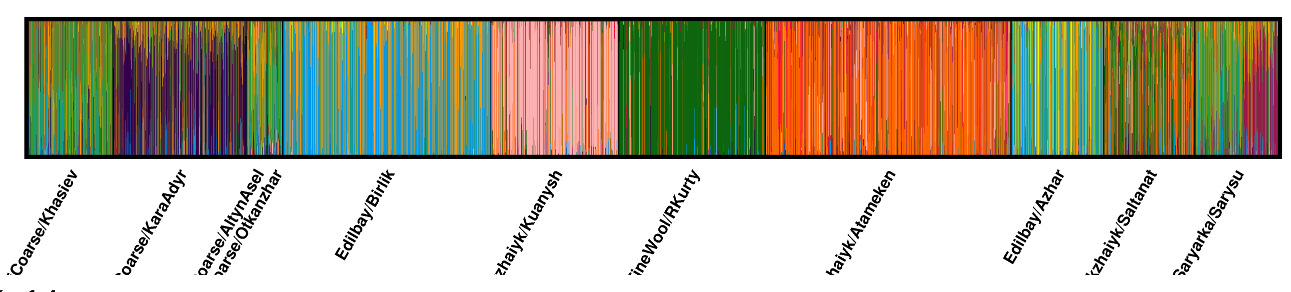

K=14

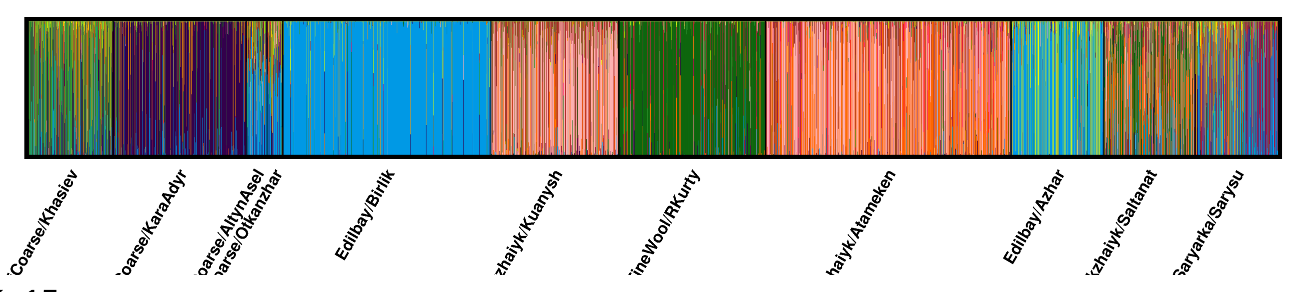

K=15

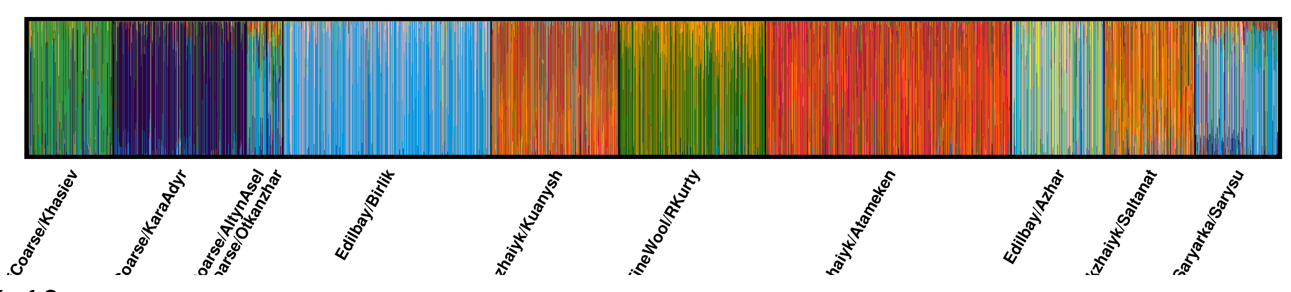

K=16

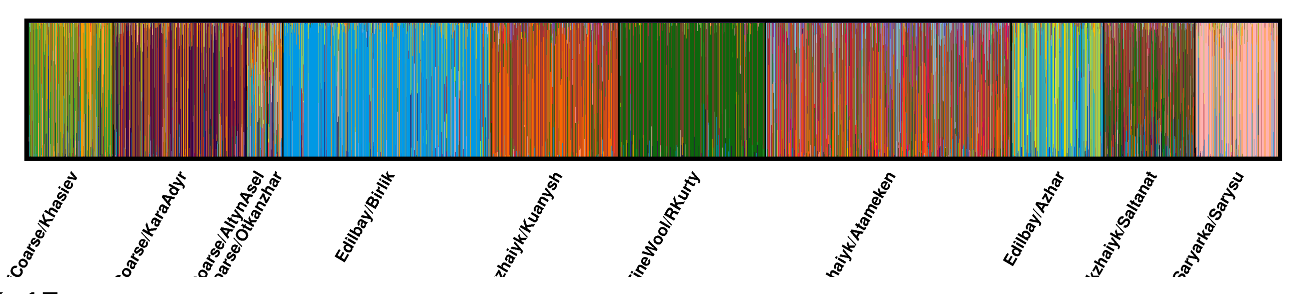

K=17

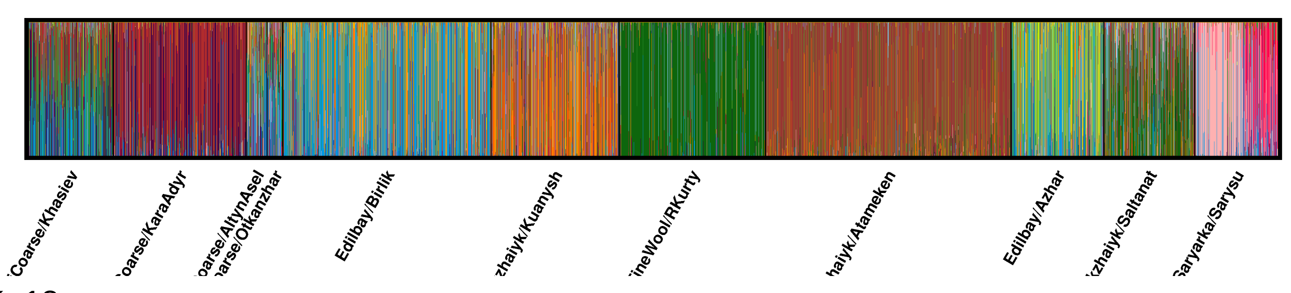

K=18

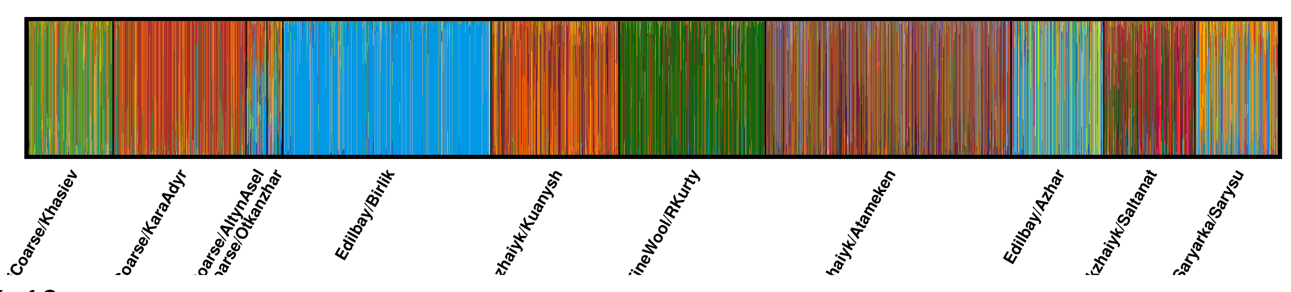

K=19

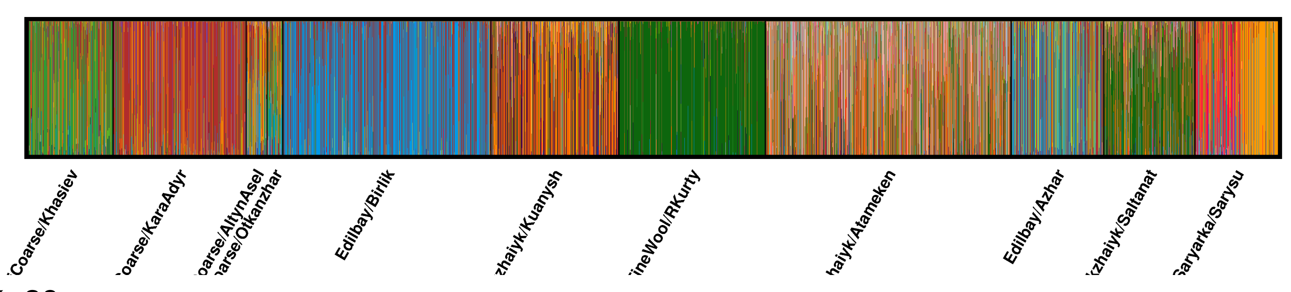

K=20

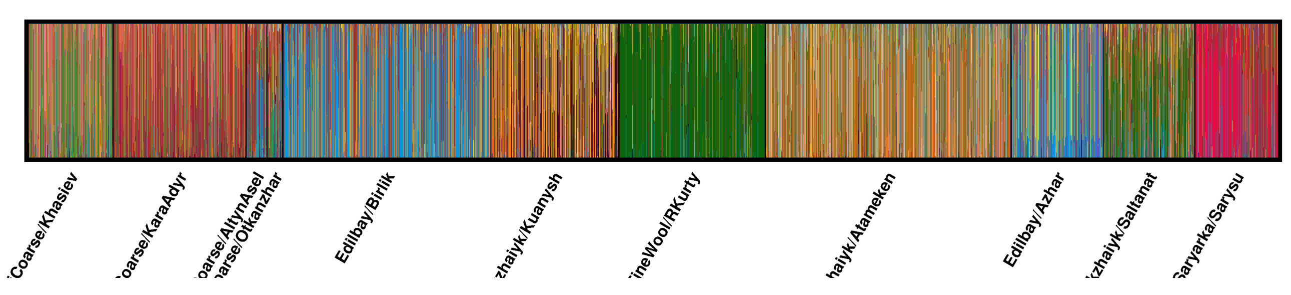

Minor modes for the uploaded data:

Division of runs by mode:

| | |
|---|---|
| K=1 | 1/1 |
| K=2 | 1/1 |
| K=3 | 1/1 |
| K=4 | 1/1 |
| K=5 | 1/1 |
| K=6 | 1/1 |
| K=7 | 1/1 |
| K=8 | 1/1 |
| K=9 | 1/1 |
| K=10 | 1/1 |
| K=11 | 1/1 |
| K=12 | 1/1 |
| K=13 | 1/1 |
| K=14 | 1/1 |
| K=15 | 1/1 |
| K=16 | 1/1 |
| K=17 | 1/1 |
| K=18 | 1/1 |
| K=19 | 1/1 |
| K=20 | 1/1 |
